# Starvation induces shrinkage of the bacterial cytoplasm

**DOI:** 10.1101/2020.12.06.413849

**Authors:** Handuo Shi, Corey S. Westfall, Jesse Kao, Pascal D. Odermatt, Spencer Cesar, Sarah Anderson, Montana Sievert, Jeremy Moore, Carlos G. Gonzalez, Lichao Zhang, Joshua E. Elias, Fred Chang, Kerwyn Casey Huang, Petra Anne Levin

## Abstract

Environmental fluctuations are a common challenge for single-celled organisms; enteric bacteria such as *Escherichia coli* experience dramatic changes in nutrient availability, pH, and temperature during their journey into and out of the host. While the effects of altered nutrient availability on gene expression and protein synthesis are well known, their impacts on cytoplasmic dynamics and cell morphology have been largely overlooked. Here, we discover that depletion of utilizable nutrients results in shrinkage of *E. coli’s* inner membrane from the cell wall. Shrinkage was accompanied by a ∼17% reduction in cytoplasmic volume and a concurrent increase in periplasmic volume. Inner membrane retraction occurred almost exclusively at the new cell pole. This phenomenon was distinct from turgor-mediated plasmolysis and independent of new transcription, translation, or canonical starvation-sensing pathways. Cytoplasmic dry-mass density increased during shrinkage, suggesting that it is driven primarily by loss of water. Shrinkage was reversible: upon a shift to nutrient-rich medium, expansion started almost immediately at a rate dependent on carbon-source quality. Robust recovery from starvation required the Tol-Pal system, highlighting the importance of envelope coupling during recovery. *Klebsiella pneumoniae* also exhibited shrinkage when shifted to carbon-free conditions, suggesting a conserved phenomenon. These findings demonstrate that even when Gram-negative bacterial growth is arrested, cell morphology and physiology are still dynamic.

**Significance statement:** Bacterial cells constantly face nutrient fluctuations in their natural environments. While previous studies have identified gene expression changes upon nutrient depletion, it is much less well known how cellular morphology and cytoplasmic properties respond to shifts in nutrient availability. Here, we discovered that switching fast-growing *Escherichia coli* cells to nutrient-free conditions results in substantial shrinkage of the inner membrane away from the cell wall, especially at the new pole. Shrinkage was primarily driven by loss of cytoplasmic water contents. Shrinkage was also exhibited by cells naturally entering stationary phase, highlighting its biological relevance across starvation conditions. The membrane-spanning Tol-Pal system was critical for robust entry into and recovery from shrinkage, indicating the importance of cell-envelope homeostasis in surviving nutrient starvation.

## Introduction

As single-celled organisms, bacteria are frequently at the mercy of their environment, with the cell envelope serving as the front line of defense. Shifts in temperature, ionic concentration, and nutrient availability result in changes in cell shape and morphology through their impacts on lipid and cell-wall synthesis and cell-cycle progression (1). Decades of work on the many ways bacteria adapt to changing environmental conditions have provided insights into transcriptional (2) and post-transcriptional (3) responses to environmental changes. Less clear is the impact of rapid changes in the environment, particularly nutrient depletion, on cell shape and physiology.

Like many gut commensals and pathogens, the model organism *Escherichia coli* experiences a dramatic change in nutrient abundance as it travels through the rich host gut into the comparatively dilute environment. Gradual depletion of nutrients is associated with exit from exponential growth and entry into the relatively quiescent state known as stationary phase (4). More rapid depletion of carbon or nitrogen leads to the starvation stringent response characterized by accumulation of the small molecules guanosine penta- and tetraphosphate (abbreviated here as ppGpp). ppGpp down regulates biosynthesis, driving exit from exponential growth, reducing metabolic activity, and eventually stimulating entry into stationary phase. Both entry into stationary phase and ppGpp accumulation have global effects on gene expression and cellular composition (5), alter the structure of the cell envelope (4), and increase resistance to diverse stresses (6).

To explore the physical effects of carbon starvation, we examined the effects of sudden removal of carbon from exponentially growing *E. coli* cells. We found that shifting from nutrient-replete to nutrient-deficient medium led to rapid growth arrest and shrinkage of the cytoplasm at the new pole in a manner distinct from osmotic shock-induced plasmolysis. We determined that shrinkage is coupled to densification of the cytoplasm and decreased abundance of translation-related proteins; shrinkage was not affected by a wide range of genetic or chemical perturbations. Cytoplasmic shrinkage also occurred during entry into stationary phase, suggesting that the root cause of shrinkage is nutrient depletion and/or an inability to access available nutrients. The rate of recovery from starvation-induced shrinkage was dependent on the quality of the re-supplied carbon source, and robust recovery required the Tol-Pal system, which is important for maintaining outer-membrane integrity. Other Gram-negative species also exhibited cytoplasmic shrinkage, suggesting that this phenomenon is a general adaptation to starvation independent of previously characterized regulatory mechanisms.

## Results

### Transition from exponential growth to carbon-free medium induces cytoplasmic shrinkage

To test the physiological effects of sudden carbon removal, we first cultured wild-type *E. coli* MG1655 cells to mid-exponential phase in LB and minimal M9 medium supplemented with 0.4% glucose (Methods). Cells sampled from each condition were washed three times in M9 salts (M9 without added carbon), resuspended in M9 salts, spotted on agarose pads containing M9 salts, and imaged via phase-contrast microscopy (Methods, Fig. 1A). Within the time required to wash and resuspend the cells and mount them on a slide (∼10 min), a readily visible phase-light region appeared at the pole of nearly every cell regardless of initial culture condition (Fig. 1B, left). In contrast to the short, stubby morphology adopted by stationary-phase *E. coli* cells (7) or in response to ppGpp accumulation (8), cells retained a rod-like shape after sudden carbon starvation (Fig. 1B, left). Importantly, resuspending cells in their original medium did not result in formation of the phase-light region (Fig. 1B, right), indicating that this phenomenon is a consequence of sudden carbon depletion. Shrinkage was not specific to M9 salts; similar behavior was observed transitioning from AB minimal salts+0.2% glucose to AB salts alone (Fig. S1A). The Gram-negative species *Klebsiella pneumoniae* also exhibited shrinkage after abrupt starvation, indicating that the phenomenon is not limited to *E. coli* (Fig. 1C). Resuspension of *E. coli* MG1655 cells cultured in M9 salts+0.4% glucose in nitrogen-free (lacking NH_4_Cl) M9+glucose similarly resulted in formation of a phase-light pole, albeit on a longer time scale (Fig. S1B), suggesting the phenomenon is generally associated with sudden depletion of nutrients.

**Figure 1:**
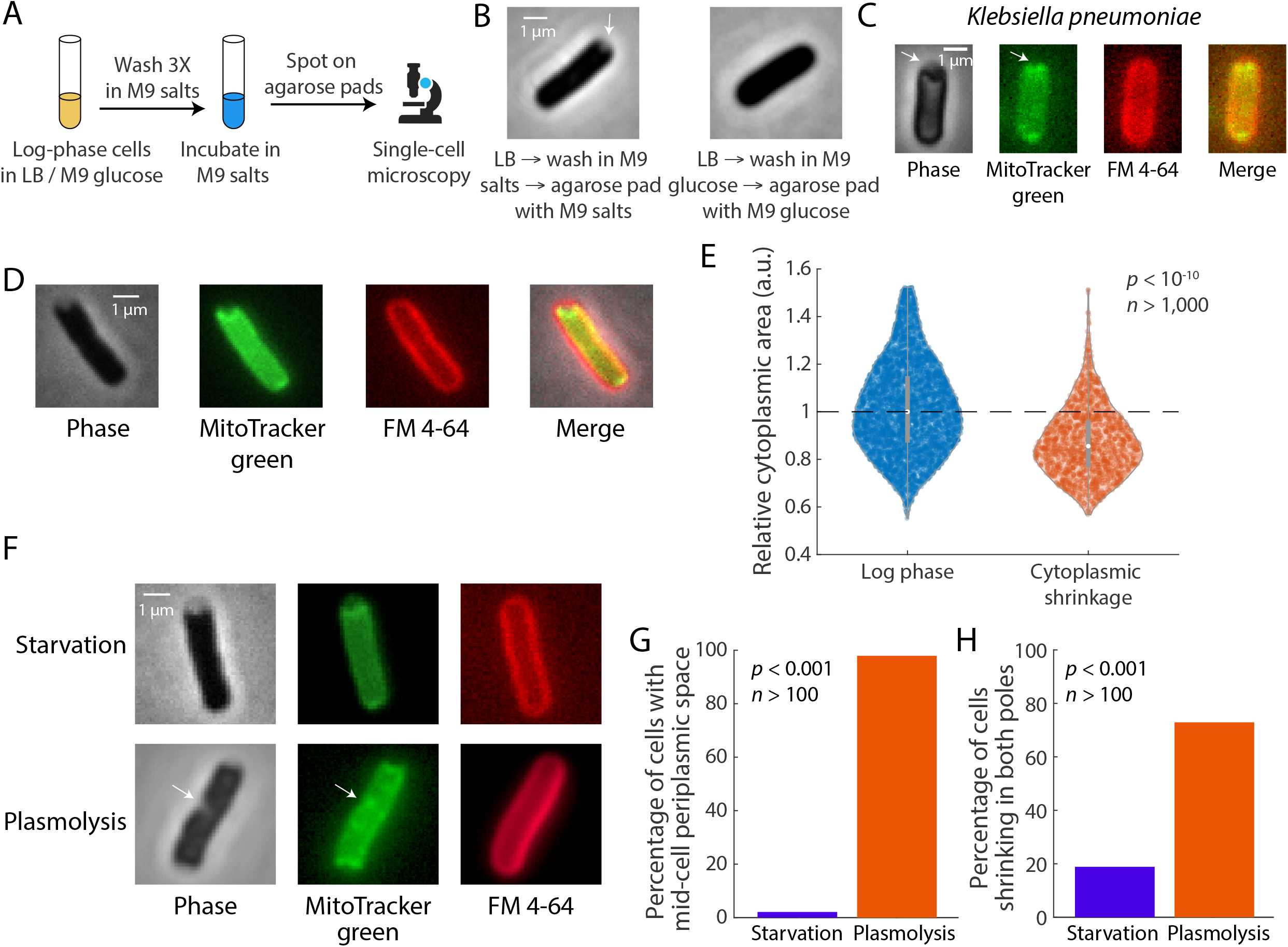
Carbon starvation causes the E. coli cytoplasm to shrink in a manner distinc from plasmolysis. A) Experimental setup for inducing cytoplasmic shrinkage. Log-phase cells were pelleted and washed in M9 salts, then incubated in M9 salts without carbon source. The resulting cells were placed onto agarose pads containing M9 salts and imaged. B) Left: log-phase cells grown in LB were washed in M9 salts three times and immediately placed on an agarose pad. Shrinkage of the cytoplasm was observed in most cells (white arrow). Right: washing in M9 glucose did not cause shrinkage. C) Shrinkage was also observed in *Klebsiella pneumoniae* cells after transitioning from LB to M9. White arrows highlight the pole with shrinkage. D) Staining of carbon-starved MG1655 cells revealed clear separation between the outer membrane dye FM4-64 and the inner membrane dye MitoTracker at one pole. E) After 4 h of starvation, the relative cytoplasmic areas of starved cells as quantified by segmenting the MitoTracker signal were ∼17% smaller than those of log-phase cells. F) Representative *E. coli* MG1655 cells 4 h after rapid starvation induced by transition from LB into M9 salts (top) or after a hyperosmotic shock with 3 M sorbitol (bottom). After hyperosmotic shock, plasmolysis bays along the cell body (arrows) were visible in phase-contrast and from MitoTracker staining, and the cytoplasm was visibly smaller at both ends. By contrast, the cytoplasm of the starved cell was shrunk only at one pole. FM4-64 stains the outer membrane, which remained largely unaffected in both cases. G, H) The percentage of cells that that exhibited mid-cell periplasmic spaces (G) or shrinkage at both poles (H) was significantly higher for osmotic shock-induced plasmolysis than for starvation (Fisher’s exact test).

The phase-light area at cell poles corresponded to an enlargement of the periplasm, the compartment between *E. coli’s* inner and outer membrane. To clarify the nature of the starvation-induced change in cell morphology, we stained MG1655 cells switched from LB to M9 salts with FM4-64 and MitoTracker, which preferentially stain the outer and inner membranes, respectively (9). Carbon starvation led to clear separation between the FM4-64 and MitoTracker contours at one end of the cell, consistent with an enlarged periplasm (Fig. 1D). We segmented the FM4-64 and MitoTracker cell contours (Methods) for log-phase and starved cells and found that starvation resulted in a ∼17% mean decrease in cytoplasmic volume (Fig. 1E).

While cytoplasmic condensation and the formation of gaps between the outer and inner membranes (plasmolysis) are characteristic of hyperosmotic shock, starvation-induced inner membrane retraction appears to be a separate phenomenon. Classically, hyperosmotic shock causes plasmolysis when the increase in external osmotic pressure is ∼100 mM (10). We measured the osmolality of LB medium to be 252 mOsm/L, and M9 salts have an osmolality of 250 mOsm/L (11). Therefore, switching from LB to M9 salts largely maintained the external osmotic pressure, and the switch from M9 glucose to M9 salts actually decreased the external osmotic pressure by ∼22 mM due to the removal of glucose. Hence, a switch to M9 salts does not induce hyperosmotic shock (12). Moreover, close examination of starved and hyperosmotic-shocked cells revealed distinct differences between the two conditions with regard to cell envelope and cytoplasmic morphology (Fig. 1F). To directly compare starvation-induced shrinkage with the effects of hyperosmotic shock, we placed MG1655 cells growing exponentially in LB on pads with LB+20% sucrose along with the inner-membrane stain MitoTracker (Methods); 20% sucrose induces an osmotic shock of ∼0.6 M. The phase-contrast and MitoTracker signals showed multiple plasmolysis bays along the body of most cells, unlike starvation-induced shrinkage (Fig. 1F,G). More than 70% of hyperosmotic-shocked cells exhibited plasmolysis at both poles, while starvation-induced shrinkage occurred predominantly at one pole (Fig. 1F,H). Thus, starvation-induced shrinkage is unlikely to be due to acute changes in the osmotic balance across the cell envelope.

### Shrinkage increases gradually with the extent and duration of nutrient removal

To better quantify periplasmic and cytoplasmic areas under various nutrient conditions, we constructed an MG1655 strain (KC1193) that expresses cytoplasmic GFP constitutively and periplasmic mCherry from the lactose promoter (Methods). We transitioned KC1193 cells from M9 salts+0.4% glucose to M9 without carbon via three washes as above, and then incubated the resuspended pellet for 4 h before imaging (Fig. 1A). Periplasmic mCherry accumulated at one pole of the overwhelming majority of cells (105/121 cells, Fig. 2A), consistent with shrinkage of the cytoplasm away from the outer membrane, unlike log-phase cells in which the mCherry signal was uniform along the cell periphery.

**Figure 2:**
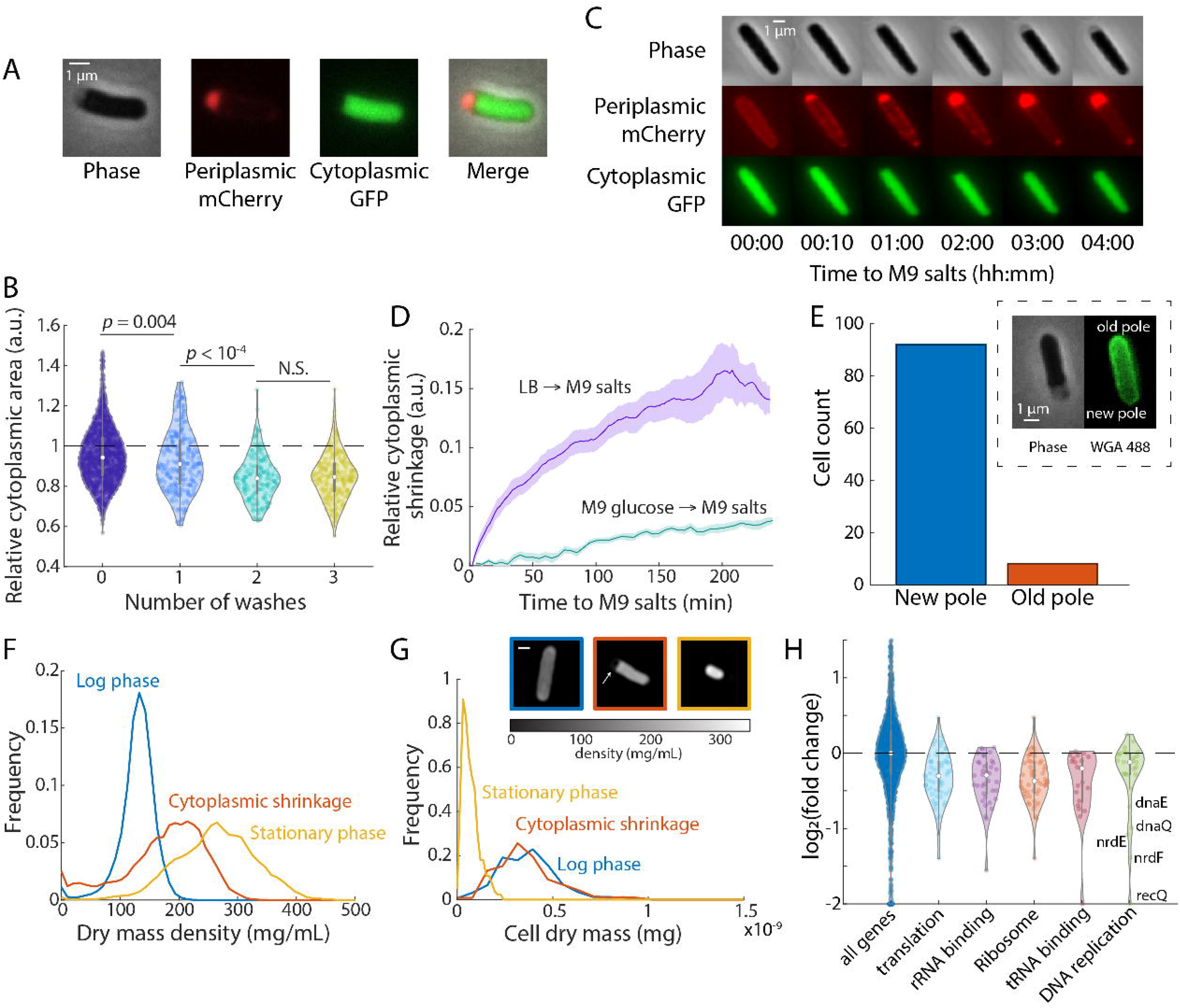
Cytoplasmic shrinkage is sensitive to the extent and duration of nutrient removal, and increases cell dry-mass density. A) Representative images of phase-contrast, cytoplasmic GFP, and periplasmic mCherry images of KC1193 cells after transitioning from LB to M9 salts for 4 h. Shrinkage of the cytoplasm was evident by the bright periplasmic mCherry signal. B) The relative cytoplasmic area of KC1193 cells (as measured by the GFP signal relative to the total cell area with GFP or mCherry signal) plateaued only after multiple washes. *n* > 100 cells for each condition, and *p*-values were calculated from two-tailed Student’s *t*-tests. C) After shifting from LB to M9 salts in a microfluidic chamber, the cytoplasmic area of KC1193 cells continued to decrease for several hours. D) The rate of shrinkage was faster in cells transitioned from LB to M9 salts compared to transitioning from M9 glucose to M9 salts. Data points are mean ± standard error of mean (sem) for n>100 cells in each condition. E) Cytoplasmic shrinkage occurs predominantly at the new pole. After pulse-chase labeling with Alexa488-WGA to label the cell periphery, the pole with separation between the cytoplasm and cell wall was almost always at the unlabeled (new) cell pole. Inset: a typical cell with shrinkage in the new (dark) pole. F) Quantitative phase imaging demonstrated that starved cells have higher dry-mass density than exponentially growing cells, yet the density is still lower compared to stationary-phase cells (n > 200 cells for each condition). G) The total dry mass of starved cells was similar to that of exponentially growing cells. Inset: typical quantitative phase images for each condition, with the white arrow highlighting the pole with shrinkage. Scale bar: 1 µm. H) Proteomics measurements highlighted classes of proteins involved in translation and DNA replication with significantly lower abundance in starved cells (*p* < 0.01, FDR < 0.05 with bootstrapping). Colored circles represent data from individual proteins. White circles are mean values across all proteins in the class and error bars are 1.5 times the interquartile range.

To determine whether shrinkage required complete removal of nutrients, we imaged KC1193 cells after 4 h of incubation after 0, 1, 2, or 3 washes in M9. Shrinkage increased after each of the first two washes, and then plateaued at ∼17% (Fig. 2B). Thus, residual nutrients are sufficient to inhibit shrinkage. To track the dynamics of shrinkage, we used a microfluidic flow cell to image KC1193 cells throughout the switch from LB to M9 salts without carbon (Fig. 2C). The cytoplasm started to shrink after <10 min and continued to shrink for ∼2 h (Fig. 2D). Similar qualitative dynamics were observed for cells switched from M9+0.4% glucose to M9 salts. After normalizing for cell volume, the rate of shrinkage was faster for cells transitioning from LB compared with M9+glucose (Fig. 2D), suggesting shrinkage may reflect a lag between nutrient depletion and the subsequent downregulation of biosynthetic capacity. Thus, shrinkage is a gradual process that starts soon after starvation and extends over hours, indicating a process highly distinct from the rapid, physical response to osmotic shock.

### Shrinkage occurs at the new cell pole

At each cell division, the mother cell creates daughter cells, with one new pole representing the site of division. Given the distinctly polar nature of cytoplasmic shrinkage following sudden starvation, we wondered if shrinkage might occur more frequently at either the old or new pole. To assess this possibility, we added Alexa488-wheat germ agglutinin (WGA) to wild-type MG1655 cells during growth (Methods) to uniformly label the cell surface (13). We then washed out the fluorescent WGA and grew the cells for approximately one cell doubling, so that the new pole would be dark while old poles remained labeled due to their relative lack of growth. We switched cells from LB to M9 salts for 4 h to induce cytoplasmic shrinkage. From a comparison of phase-contrast and WGA fluorescence images, shrinkage clearly occurred at the new (unlabeled) pole in the vast majority of cells (>90%) (Fig. 2E), suggesting that attachment between the inner membrane and cell wall/outer membrane is weaker at the new pole than at the old pole.

### Starvation-induced cytoplasmic shrinkage is characterized by increased cell density

To determine whether cells lose material during shrinkage, we assessed dry-mass cell density before and after shrinkage via quantitative phase imaging, which uses *z*-stacks of brightfield images to compute density-dependent phase shifts (14–18). Consistent with previous reports (14, 15), dry-mass density in the cytoplasm was ∼80% higher in stationary phase than in log phase (Fig. 2F). Highlighting that the shrunken state is distinct from stationary phase, the dry-mass density of suddenly starved cells (transitioned from LB to M9 salts for 4 h) was intermediate between the densities of stationary- and log-phase cells (Fig. 2F). Importantly, the density increase in starved cells relative to log-phase cells was ∼25%. Considering that the cytoplasmic volume shrunk by ∼17%, the density increase largely offsets the decrease in cytoplasmic volume and maintains total cell dry mass (relative mass of the shrunken cells = (1+0.25) x (1.04). Indeed, the distribution of total dry mass per cell was highly similar between log-phase and starved cells (Fig. 2G). The periplasmic region remained at a dry-mass density close to zero. Thus, these results suggest that cells shrink by losing water while retaining most of their cytoplasmic contents.

### Shrinkage is not coupled to a change in cytoplasmic diffusion

Based on increases in cytoplasmic density (Fig. 2F), we wondered whether diffusion was limited in starved cells. Reduced rates of cytoplasmic diffusion have been observed in *E. coli* cells after hyperosmotic shock-induced plasmolysis (19) and treatment with 2,4-dinitrophenol (DNP) (20), an oxidative phosphorylation uncoupler that depletes cells of ATP energy. To assess cytoplasmic properties after sudden nutrient depletion, we measured the diffusion constant of a GFP-labeled self-assembling viral protein µNS (20). The larger size of the assembled µNS particles results in slower diffusion than single proteins, making their motion easier to track than the motion of individual GFP molecules. We fit the slope of mean-squared displacement over time to extract a diffusion constant along the length dimension defined by the cell midline (Fig. S2A, Methods), and our measured diffusion constants for exponentially-growing cells were consistent with previous results (20). Somewhat surprisingly, while DNP treatment reduced diffusion as expected (20), the diffusion constant was essentially equivalent in exponentially growing cells and in starved, shrunken cells (Fig. S2B). These findings indicate that the increase in cytoplasmic density during acute carbon starvation is insufficient to alter the diffusive motion of these nanoparticles, and suggest that the impact of starvation on metabolic activity is distinct to that of DNP treatment.

### Shrinkage is independent of biosynthesis, stress-response pathways, and regulatory circuits governing cell-envelope homeostasis

Precisely orchestrated changes in transcription, translation, and/or proteolysis are characteristic of diverse bacterial stress-response pathways. Inhibiting any one of these major biosynthetic processes typically interferes with the ability of cells to adapt to new conditions. To determine whether new biosynthesis is required for cytoplasmic shrinkage, we added the following antibiotics to M9 salts during the initial washes and during resuspension: rifampicin, to inhibit transcription; chloramphenicol, to inhibit translation; cerulenin, to inhibit lipid synthesis; and A22, to inhibit cell-wall synthesis. In all cases, shrinkage still occurred, indicating that the phenomenon is independent of major biosynthetic pathways (Fig. S2C).

Next, we sought to determine the effect of previously characterized stress-response pathways associated with nutrient limitation on shrinkage. In *E. coli*, depletion of carbon and/or nitrogen induces accumulation of ppGpp, resulting in down-regulation of a wide range of biosynthetic pathways and a concomitant reduction in growth (21, 22). The transcriptional regulator RpoS (SigmaS), which is induced in response to diverse stresses (23, 24), is also important for entry into stationary phase, a state associated with nutrient depletion as well as cell crowding. Surprisingly given the central roles of RpoS and ppGpp in starvation-induced stress responses, cells defective in RpoS (Δ*rpoS*) or ppGpp synthesis and accumulation (Δ*relA* Δ*spoT*) exhibited cytoplasmic shrinkage after washing and incubation in M9 salts (Fig. S2D). Δ*rpoS* cells were phenotypically indistinguishable from wild-type cells in this regard. Populations of ppGpp mutants are notoriously heterogeneous in size, with ∼10% of cells forming long aseptate filaments (25). Nevertheless, periplasmic expansion was readily visible at the poles of Δ*relA* Δ*spoT* cells regardless of the degree of filamentation (Figure S2D), indicating that shrinkage was independent of ppGpp production. Shrinkage also occurred in cells lacking ClpP (Figure S2D), a serine protease required for adaptation to and extended viability in stationary phase (26, 27). Thus, as with the chemical inhibitors of biosynthesis described above, the degree and frequency of shrinkage was independent of all of these stress-response pathways.

Starvation-induced shrinkage also occurred in cells defective in carbon acquisition and metabolism. Deletion of the gene encoding Crr, a phosphotransfer protein involved in the uptake and phosphorylation of glucose and several other carbon sources (28), or Pgi, which catalyzes the interconversion of glucose-6-phosphate and fructose-6-phosphate during glycolysis and gluconeogenesis (29), had no impact on the timing or extent of cytoplasmic shrinkage (Fig. S2D). Finally, despite a superficial similarity between shrunken cells and cells defective in outer membrane lipid homeostasis (7), deletion of MlaA, a lipoprotein involved in maintenance of outer-membrane lipid asymmetry through retrograde phospholipid trafficking (30), did not affect the frequency or degree of shrinkage (Fig. S2D). Thus, shrinkage does not require biosynthesis, induction of known stress-response pathways, or regulation of cell-envelope homeostasis.

### Proteins involved in translation and DNA replication are down-regulated during starvation

To determine if starvation-induced shrinkage is associated with distinct shifts in proteome composition, we extracted proteins from log-phase cells and cells acutely starved for 4 h, and analyzed protein abundances using mass spectrometry (Methods, Table S1). No protein classes exhibited a significantly higher protein abundance in starved cells versus log-phase cells, and the only gene classes with a significant decrease in starved cells involved translation or DNA replication (Fig. 2H); for the latter, the slight ∼5% average decrease was predominantly due to lower levels of five proteins (Fig. 2H). Thus, only the machinery responsible for rapid growth is affected by acute starvation.

### The rate of cytoplasmic recovery depends on the quality of carbon source

Importantly, shrinkage is reversible in wild type cells. Plating wild-type MG1655 and BW25113 cells 4 h after a shift from LB to M9 salts resulted in ∼100% recovery (Fig. 3A). The viability of shrunken cells was confirmed by staining them with propidium iodide (Fig. 3B), a DNA intercalator that only enters cells with compromised plasma membranes and as such is frequently used as a live/dead indicator (31).

**Figure 3:**
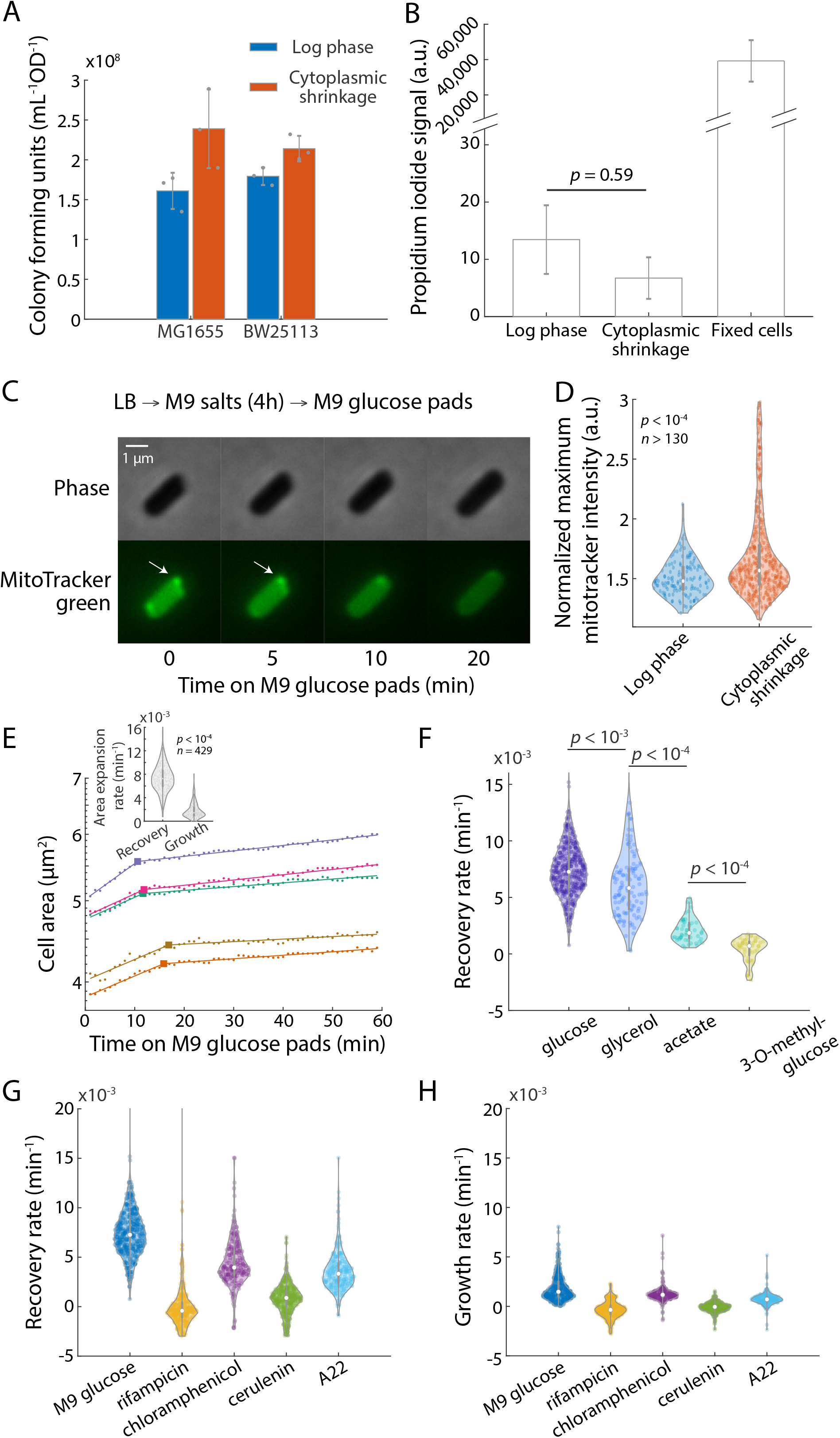
Recovery from starvation is slow and dependent on carbon-source quality. A) Starvation did not reduce viable cell counts in either MG1655 and BW25113 cells, suggesting that all cells recovered from shrinkage and were able to regrow when exposed to fresh nutrient. Data points are mean ± standard deviation (S.D.) with *n*=3 replicates. B) Propidium iodide staining was similarly low (*n*>100 cells, two-tailed Student’s *t*-test) in exponentially growing and starved cells as compared with cells killed and fixed with ethanol. C) Cells starved in M9 salts for 4 h were spotted onto an agarose pad with M9+0.4% glucose and MitoTracker and monitored using time-lapse microscopy. The cytoplasm began to re-expand almost immediately after cells were placed on the agarose pad. Initially, cells exhibited bright MitoTracker foci, suggesting membrane crumpling, and the foci appeared to dissolve as the cytoplasm re-expanded. D) MitoTracker signal is more uniformly distributed in exponentially growing cells compared with starved cells (two-tailed Student’s *t*-test), consistent with the observation of inner membrane puncta in starved cells. E) Typical single-cell traces of cytoplasmic area expansion after starved cells were exposed to fresh nutrient again. The expansion rate was faster during the recovery phase when inner membrane was shrunk away from the cell wall, but slowed down after the inner membrane rejoined the cell wall. Individual round dots are raw data, solid lines are best fits to exponential curves, and squares are the best fit for the transition time from fast to slow expansion. Inset: area expansion rate during recovery was faster than growth post-recovery (two-tailed Student’s *t*-test). F) The recovery rate after starvation was dependent on the quality of carbon sources in the recovery media. Recovery was slower on carbon sources that support slower steady-state growth rates. Addition of the non-metabolizable glucose analog 3-O-methyl-glucose did not stimulate recovery. *n*>100 cells for each condition. *p*-values are from two-tailed Student’s *t*-tests. G,H) In the presence of different antibiotics, starved cells still recovered when exposed to fresh glucose. The recovery (G) and growth (H) rates after starvation were slower (*p*<10^−4^ in all conditions, Student’s *t*-test, *n*>100 cells) compared to the antibiotic-free control. In (D), (F), (G), and (H), colored circles represent data from individual cells. White circles are mean values across all cells in that condition and error bars are 1.5 times the interquartile range

To determine the dynamics of recovery, we placed cells starved in M9 salts onto an M9+0.4% glucose agarose pad with MitoTracker stain and monitored regrowth using time-lapse microscopy (Methods). The cytoplasm began to re-expand almost immediately after the cells were placed on the agarose pad (Fig. 3C), indicating that the cells were ready to regrow despite prolonged starvation. Notably, starved cells exhibited bright foci of MitoTracker signal at the boundary of the cytoplasm (Fig. 3C, white arrows); such foci are indicative of folded membrane (32) and suggest that shrinkage is not due to cells metabolizing membrane components and thereby compressing the cytoplasm. In support of this interpretation, the maximum MitoTracker signal after normalization was significantly higher in starved cells than in log-phase cells (Fig. 3D). Plasma-membrane foci dissolved as the cytoplasm re-expanded, with complete dissolution coinciding with the cytoplasm rejoining the cell wall (Fig. 3C).

We further quantified the rate of normalized area expansion during recovery. Expansion was initially relatively fast (0.007 min^−1^), but once the cytoplasm reached the size of the cell envelope, expansion slowed down to <0.002 min^−1^ (corresponding to a doubling time of >5 h, Fig. 3E). We term these two expansion phases as the “recovery” and “growth” phases, respectively. Even during the faster recovery phase, the rate of expansion was slower than the growth rate of exponentially growing cells in M9+0.4% glucose (∼0.01 min^−1^), consistent with a reduction in translational machinery (Fig. 2H). The rapid slowdown in growth once the cytoplasm reached the size of the rest of the cell envelope (Fig. 3E) suggests that the cell wall either mechanically resisted expansion or that growth of the cell envelope was slower than cytoplasmic growth during the period over which the cells were tracked.

To determine whether recovery was dependent on the presence of a metabolizable carbon source, we examined the behavior of starved cells on M9 agarose pads supplemented with a variety of carbon sources. The initial recovery rate was slower on carbon sources that support slower steady-state growth rates (Fig. 3F). Importantly, addition of 3-O-methyl-glucose, which is imported and phosphorylated rather than metabolized (33), did not stimulate recovery (Fig. 3F), indicating that recovery requires synthesis. These data, along with the observation that the first recovery phase requires tens of minutes (Fig. 3E) rather than the seconds required to swell a cell after hypoosmotic shock (10), indicate that metabolism drives recovery.

To probe potential drivers of recovery after cytoplasmic shrinkage, we examined the impact of antibiotic treatment on recovery. Cytoplasmic re-expansion was observed in the presence of rifampicin, chloramphenicol, cerulenin, and A22, although all four compounds slowed down recovery and outgrowth (Fig. 3G,H). Together, these data suggest that while re-expansion relies on active metabolism of fresh nutrients, it does not directly depend on new transcription, translation, or envelope synthesis.

### Analysis of stationary-phase cells suggests that shrinkage is a general response to depletion of useful nutrients

To determine whether cytoplasmic shrinkage constitutes a more general response to nutrient depletion beyond abrupt starvation, we examined stationary-phase KC1193 cells after 18 h of growth in LB. In these cells, periplasmic mCherry was clearly concentrated at one pole, signifying some degree of cytoplasmic shrinkage (Fig. 4A). Surprisingly, shrinkage was visible in some cells in phase-contrast images (Fig. 4A, white arrows), suggesting that it had been overlooked in previous studies of stationary-phase cell morphology.

**Figure 4:**
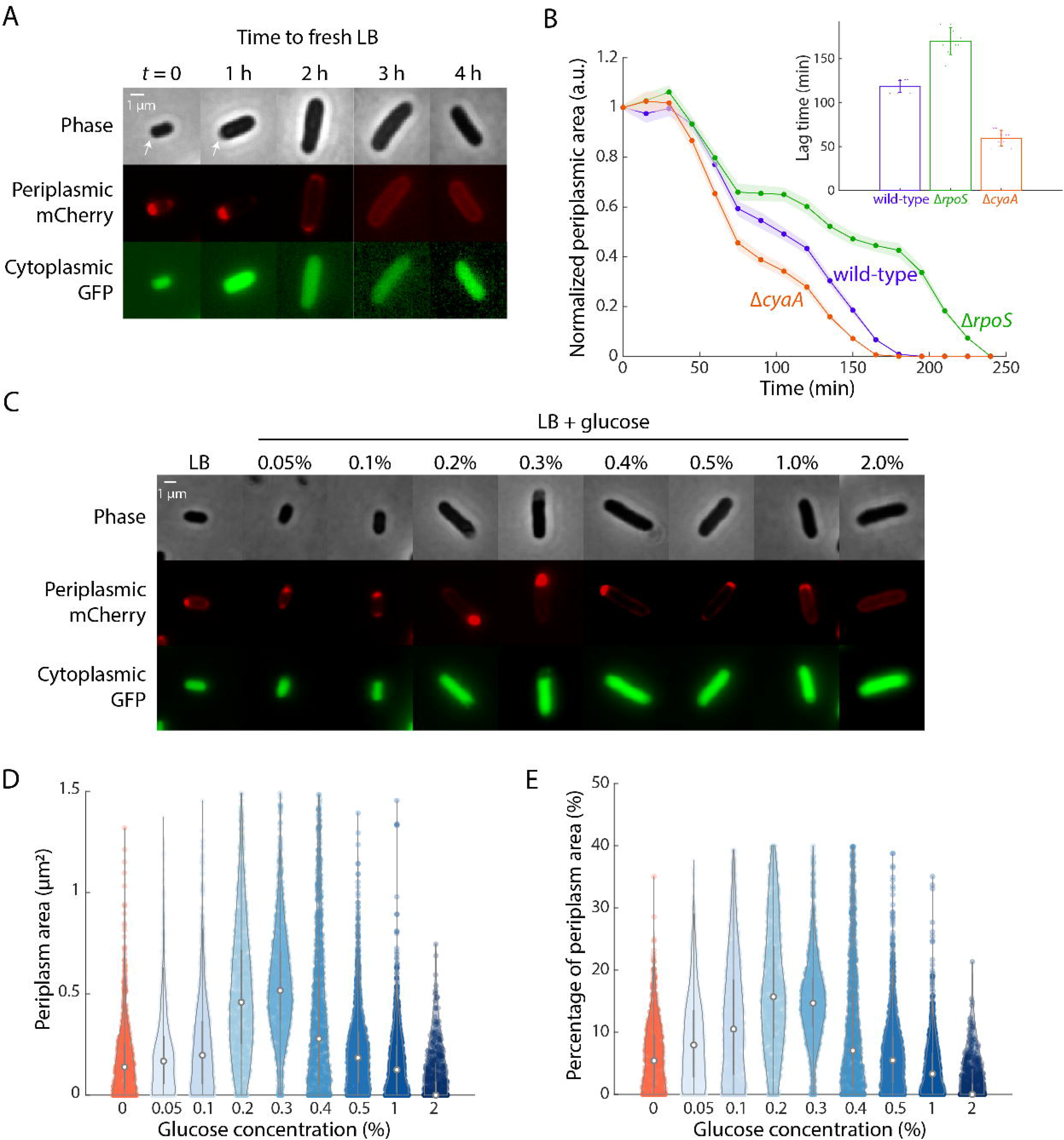
Shrinkage occurs during stationary phase and is exacerbated by intermediate levels of glucose. A) Stationary-phase KC1193 cells exhibited a large periplasmic region that decreased in area over several hours after the cells were incubated in fresh LB. B) The periplasmic area in Δ*rpoS* and Δ*cyaA* cells expressing periplasmic mCherry resolved slower and faster, respectively, than in wild-type cells, consistent with their differences in lag time (inset). Data points are mean ± sem with *n*>100 single cells. Inset: lag times of each strain. Data points are mean ± S.D. with *n*=8 replicates. C-E) Cells grown in LB+0.2-0.3% glucose have larger periplasmic spaces than cells grown in LB or LB+2% glucose. Images in (C) are of KC1193 cells. The total periplasmic area (D) and the periplasmic area normalized to total cell area (E) are highest in LB+0.3% glucose. *n*>100 cells for each condition.

Resuspension of stationary-phase KC1193 cells in fresh LB resulted in resolution of polar mCherry signal within ∼150 min (Fig. 4A,B, S3A), a time frame similar to the population based-lag time determined by optical density (Fig. 4B, inset). Mutants that increased (e.g. Δ*rpoS*) or reduced (Δ*cyaA*) lag time resolved periplasmic area in a longer and shorter time period, respectively (Fig. 4B, S3A), consistent with resolution reflecting the ability of different strains to ramp up metabolism. Shrinkage was also observed in KC1193 cells cultured overnight in LB supplemented with a range of glucose concentrations from 0-2%: cells grown overnight in LB+0.2-0.3% glucose exhibited the most obvious shrinkage, both in total periplasmic area (Fig. 4C) and relative to cell size (Fig. 4D). Another study also reported that abrupt nutrient down-shift or glucose starvation of *E. coli* cells led to slight shrinkage in the cytoplasmic volume (14), consistent with our observation that shrinkage is connected to nutrient depletion. Thus, shrinkage is a general, sometimes dramatic, feature of stationary-phase as well as abruptly starved cells, and the extent of shrinkage varies across mutants and nutrient conditions.

### Shrinkage is directly connected with growth during starvation and recovery

To assess the kinetics of shrinkage in relation to cell growth, we shifted log-phase KC1193 cells from fresh LB to spent medium (filtered supernatant from an 18-h stationary-phase culture) and back to fresh LB after 100 min in a microfluidic flow cell (Fig. 5A, Methods). We employed spent LB rather than AB salts to ensure that osmolarity remained constant throughout the experiment and to further interrogate the nature of shrinkage during stationary phase. Growth was immediately curtailed following a switch to spent LB and halted completely within 10 min (Fig. 5B). We detected increases in periplasmic area only after growth completely ceased (Fig. 5B), further supporting the conclusion that shrinkage only occurs in the absence of cell growth. The addition of fresh LB at *t=*110 min resulted in an immediate increase in both cytoplasmic volume and growth rate (Fig. 5B), suggesting that recovery from shrinkage is connected with growth resumption under these conditions. As with recovery from abrupt starvation, the gradual increase in growth rate is consistent with the decrease in abundance of translation-related proteins during shrinkage (Fig. 2H).

**Figure 5:**
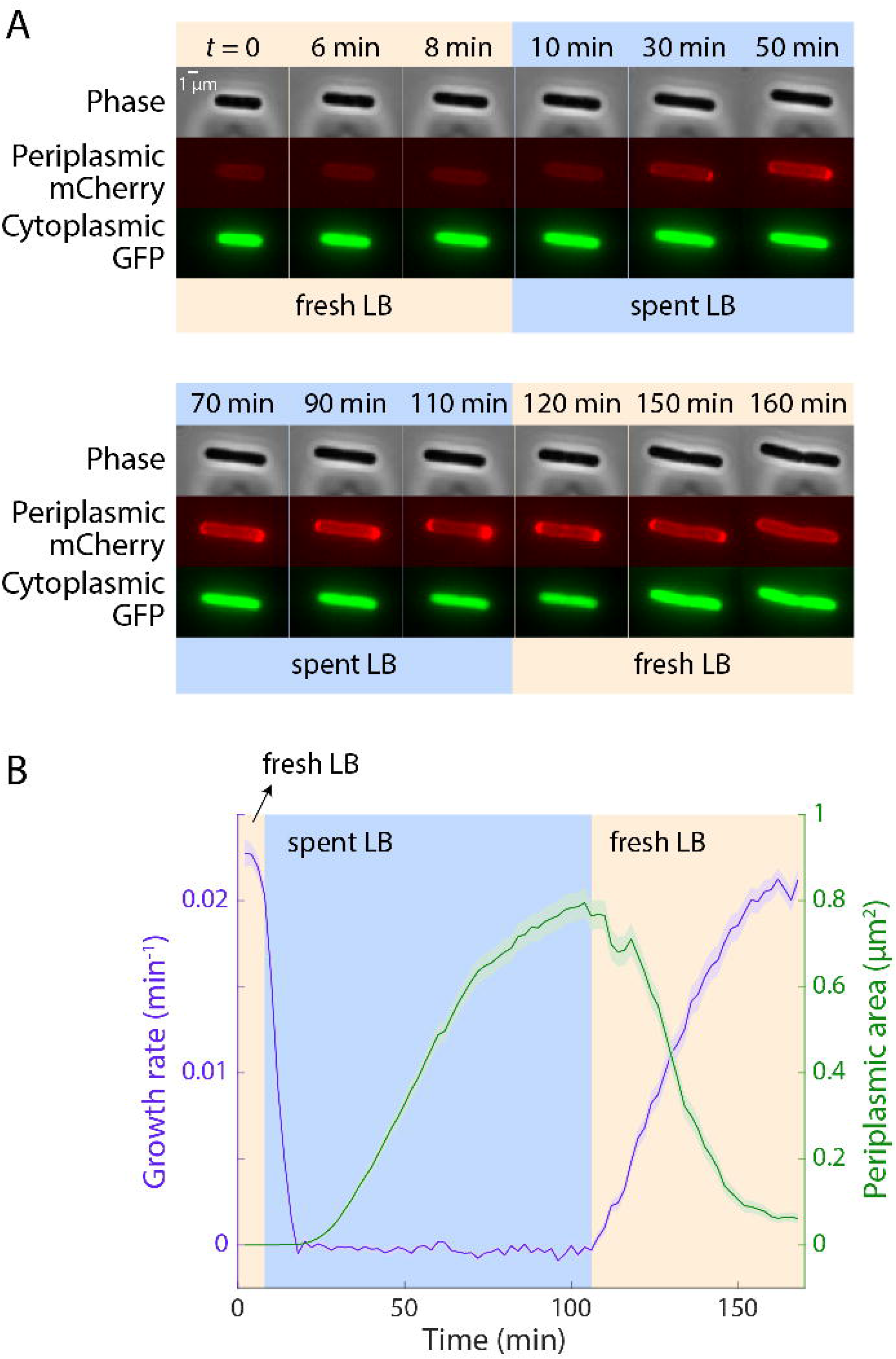
Shrinkage initiates only once growth rate drops to zero, while gradual growth resumption and resolution of periplasmic space occur simultaneously. A) Time-lapse images of KC1193 cells grown in fresh LB in a microfluidic flow cell for 10 min, then switched to spent LB for 100 min, then switched back to fresh LB. The osmolalities of the fresh and spent LB were equalized. The periplasmic area has expanded after 20 min in spent LB, and gradually shrinks after the resumption of growth in fresh LB. B) Quantification of single-cell growth rate and periplasmic area revealed that growth dropped to zero over ∼10 min after switching to spent LB. The periplasmic area only started to increase after the growth rate reached zero, and increased steadily over the 100 min in spent LB. After the switch back to fresh LB, growth rate immediately started to increase, gradually reaching pre-spent LB levels after >60 min. The resolution of the periplasmic area was coincident with the growth-rate increase. Data points are mean ± sem with *n*>150 cells.

### A functional Tol-Pal system is required for shrinkage and recovery

Although disruption of cell-wall synthesis during elongation via A22 treatment did not alter shrinkage or recovery (Fig. 3G,H, S2C), the predominance of shrinkage at the new pole (Fig. 2E) led us to hypothesize that disruption of cell division would alter shrinkage or recovery. The Tol-Pal system plays a role in outer-membrane invagination and septal peptidoglycan processing during cell division (34) and is important for maintaining outer membrane integrity (35). Intriguingly, carbon-starved Δ*tolA* cells exhibited diverse single-cell phenotypes (Fig. 6A): a small subpopulation exhibited shrinkage like wild-type cells (36/209), while the majority of cells either lysed (62/209) or developed membrane blebs at cell poles or mid-cell (78/209), and another subpopulation exhibited chaining (16/209). Similar phenotypes were also observed for other genetic deletions in the Tol-Pal pathway, including Δ*tolB*, Δ*tolR*, and Δ*pal* (Fig. S4). All Tol-Pal pathway deletion mutants also exhibited 50-70% lower CFU yields after starvation-induced shrinkage (Fig. 6B).

**Figure 6:**
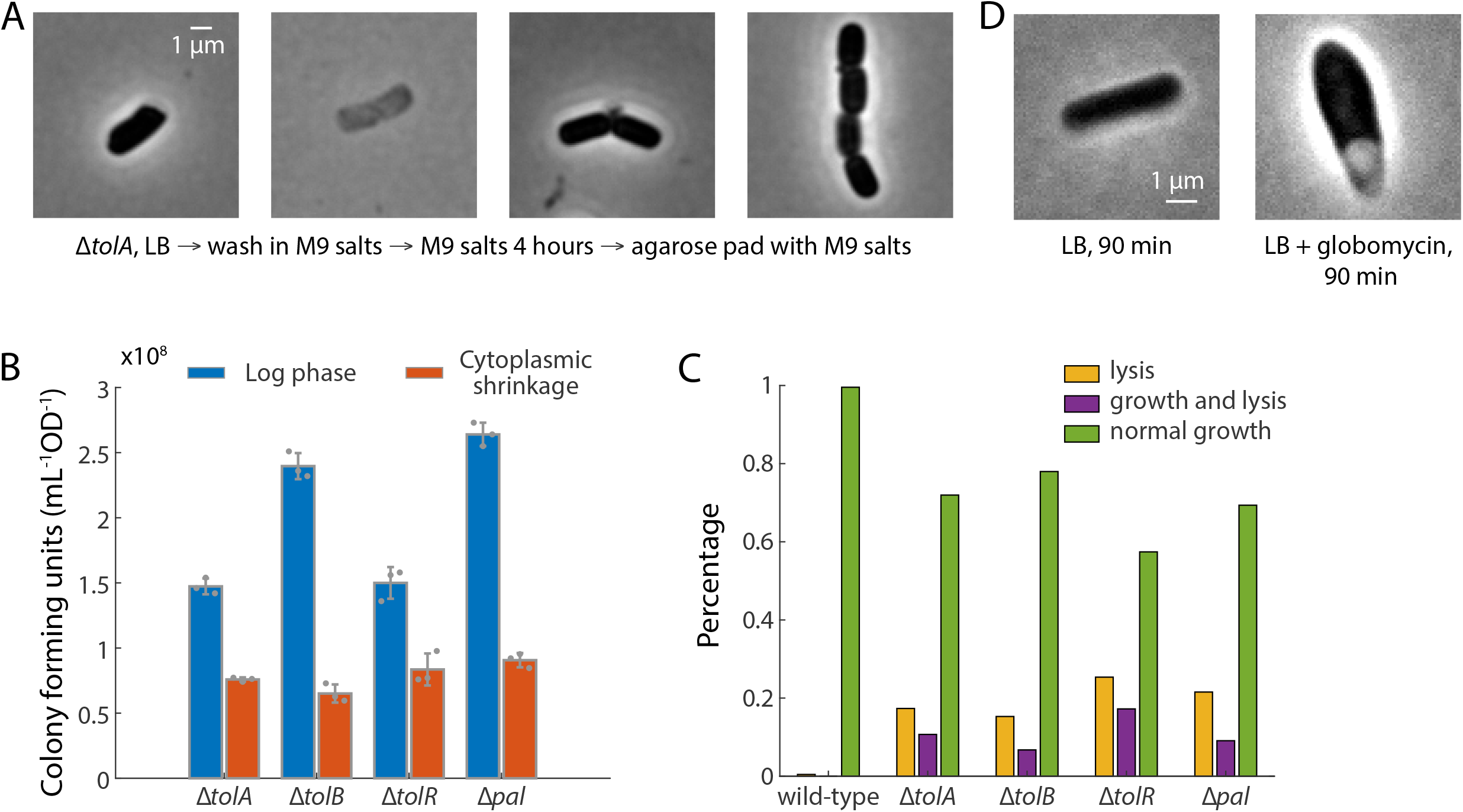
Shrinkage and recovery are dependent on a functional Tol-Pal system. A) After 4 h of starvation in M9 salts, Δ*tolA* cells exhibited diverse phenotypes including shrinkage, lysis, blebbing and chaining. B) The Tol-Pal pathway maintains viability during recovery from sudden starvation. All deletion mutants in the Tol-Pal pathway had 50-70% lower CFU yield compared to exponential phase, by contrast to the wild-type for which CFU levels were maintained (Fig. 3A). Data points are mean ± standard deviation (S.D.) with *n* = 3 replicates. C) Quantification of the fraction of cells that outgrew from stationary phase. While virtually all wild-type cells resumed growth, 20-40% of Tol-Pal mutant cells exhibited lysis within 3 h, suggesting that the Tol-Pal pathway is also critical during recovery from normal stationary phase. D) Treatment of exponentially growing cells with the lipoprotein signal peptidase II inhibitor globomycin causes shrinkage. MG1655 cells were treated with 20 µg/mL globomycin for 90 min.

Since the shrinkage phenotype was also induced naturally in stationary-phase cells (Fig. 4A), we further investigated the Tol-Pal deletions in stationary phase. The morphologies of stationary-phase Δ*tolA* cells were similar to wild type. However, when Δ*tolA* cells were placed onto agarose pads containing fresh medium, only a fraction resumed growth (unlike wild-type cells in which virtually all cells regrew): during a 3-h time-lapse imaging of 271 Δ*tolA* cells, 47 cells exhibited no growth and eventually lysed, 29 cells first grew but then lysed before the first cell division, and only 195 cells resumed growth normally (Fig. 6C). Similar trends were observed in other Tol-Pal deletions (Fig. 6C). Taken together, these data suggest that coordination between the inner and outer membranes is essential for efficient recovery from starvation.

To test the hypothesis that shrinkage results from a disruption of coordination between the inner and outer membranes, we were motivated by previous studies showing that treatment with globomycin, an inhibitor of outer membrane biogenesis through its impact on activity of lipoprotein-specific signal peptidase II (LspA), induces condensation of the bacterial chromosome (36). We tested the impact of 20 ug/mL globomycin (∼2X MIC) on the morphology of exponentially growing wild-type cells. Globomycin treatment resulted in a phenotype nearly identical to that observed in carbon-starved cells, with a reduction in cytoplasmic volume and enlargement of the periplasm at a single pole (Fig. 6D), supporting disruption of inner and outer membrane biogenesis as a driver of shrinkage.

## Discussion

In this study, we determined that two types of starvation—sudden limitation of nutrients and stationary phase—induces the *E. coli* cytoplasm to shrink away from the cell wall and outer membrane (Fig. 1), increasing cytoplasmic density ∼1.3-fold (Fig. 2F). Our data suggest that homeostatic connections between inner and outer membrane biogenesis play a role in both shrinkage and recovery. Treatment with globomycin, an inhibitor of lipoprotein secretion and outer membrane biogenesis, induced shrinkage in exponentially growing cells that was phenotypically indistinguishable from that caused by sudden nutrient depletion (Fig. 6D). Consistent with the notion that defects in inner and outer membrane coordination contribute to shrinkage, a gain-of-function mutation in MlaA (*mlaA\**), which contributes to retrograde phospholipid transport (30), leads to cytoplasmic shrinkage when *E. coli* is resuspended in spent LB medium (7). The lipoprotein MlaA removes phospholipids from the outer leaflet of the outer membrane and shuttles them to the periplasm, making it important for outer membrane asymmetry (30). Shrinkage in exponentially growing cells with altered phospholipid synthesis (37) further indicates that dysbiosis between inner and outer membrane biogenesis is a major contributor to shrinkage.

Recovery from cytoplasmic shrinkage appears to be similarly dependent on inner and outer membrane coordination. Although they do not appear to impact shrinkage, defects in the Tol-Pal system, periplasmic proteins implicated in linkages between inner membrane, cell wall and outer membrane, severely impaired recovery of stationary-phase cells. Stationary phase Δ*tolA* mutants exhibited blebbing and lysis upon resuspension in fresh medium, a fate at least superficially similar to the demise of *mlaA\** mutants after an extended period in spent LB (7).

Together, these results suggest a model in which loss of connections between outer and inner membrane synthesis contribute to shrinkage and that restoration of these connections is critical to recovery. The strong bias toward sudden nutrient starvation-induced shrinkage at the new pole (Fig. 2E) suggests that connections between the inner and outer membrane develop subsequent to division, making the new pole more vulnerable in this regard. The contribution of linkages between inner and outer membrane to shrinkage and recovery is further supported by the importance of the Tol-Pal system to efficient recovery from stationary phase (Fig. 6D). Previously implicated in maintaining viability during division (34), Tol-Pal may preserve cell envelope integrity in starved and stationary-phase cells by maintaining connections between the inner and outer membranes as the inner membrane shrinks and expands. It remains unclear whether loss of connections between the membranes directly causes water loss from the cytoplasm and hence condensation. *E. coli* cells were previously shown to shrink substantially after a week of starvation, although some of this shrinkage was due to loss of cellular material into the extracellular milieu (38). Regardless, observations of starvation-induced cytoplasm shrinkage in *K. pneumoniae* (Fig. 1C) and in *Vibrio cholerae* (39) suggest that shrinkage may be widespread across Gram-negative bacteria.

Given the potentially generic nature of shrinkage across starvation conditions and the independence of a variety of pathways, one possibility for the mechanism underlying shrinkage is that the condensed cytoplasm is a passive result of the lack of metabolic activity, akin to a previous hypothesis that the cytoplasm enters a “glassy” (less fluid) state with dramatically slowed diffusion in energy-depleted conditions (20). However, µNS diffusion was unaffected by starvation and shrinkage (Fig. S2A,B), suggesting that energy depletion via starvation impacts cell physiology considerably differently than energy depletion induced by an uncoupler. We previously discovered that diffusivity does not change dramatically throughout density changes in fission yeast induced by osmotic oscillations (16), suggesting that increased cytoplasmic density and/or reduced metabolic capacity need not inhibit fluidization.

It is possible that shrinkage as cells lose water (Fig. 2F,G) is caused by a gradual decrease in turgor pressure; a recent study found that immediately after nutrient upshift volume increased more quickly than dry mass, similar to a hypoosmotic shock (14). However, the mechanism by which cells would achieve such osmoregulation has yet to be determined; an intriguing speculation motivated by the defects in Tol-Pal mutants is that the cell wall produces compressive forces that exert osmotic-like effects.

The extent to which starvation and shrinking can be beneficial (or even tolerated) is unclear. Is cytoplasmic densification an adaptive benefit or a passive response to starvation? One possibility is that carbon depletion stimulates cells to shrink as a way to increase the concentration of the metabolic intermediates and enzymes involved in carbon metabolism, thereby driving reactions forward. This behavior would allow bacteria to turn ‘worthless’ intermediates into useful products/energy. Such a mechanism would be similar to how enzyme clustering accelerates the processing of intermediates through metabolic channeling (40), and is consistent with the tight coupling between shrinkage and growth rate as cells emerge from stationary phase (Fig. 5). Identification of a mutant that does not exhibit detachment would empower metabolomics studies to test this hypothesis, but we have been unsuccessful in identifying such a mutant using targeted approaches (Fig. S2). There is likely a maximum interval of starvation that *E. coli* cells can generally withstand, as cell viability eventually drops during development of the GASP phenotype (41). Protein degradation may also eventually lead to cells not being able to sense or use nutrients to expand the inner membrane back to the cell wall, particularly since translation-related proteins experience the largest decrease during starvation detected here (Fig. 2H). In budding yeast, macromolecular mobility is restricted by glucose starvation due to a reduction in cell volume (42) and by chemical inhibition of glycolysis (43). In the future, it will be interesting to determine the generality of this behavior across organisms; the presence or absence of shrinkage may reflect different environmental pressures during starvation. Understanding the response of bacterial cells to rapid removal of nutrients may also indicate mechanisms for surviving stressful environments. Ultimately, the discovery of novel starvation responses in bacteria may provide insights into the dynamics of host-host transmission, particularly in pathogens, and reveal novel targets such as Tol-Pal for disrupting starvation recovery.

## Author Contributions

H.S., C.W., K.C.H., and P.L. conceived of and designed the research; H.S., C.W., J.K., P.D.O., S.C., S.A., M.S., J.M., C.G.G., and L.Z. performed the research; H.S., C.W., J.K., K.C.H., and P.A.L. analyzed the data; and H.S., C.W., F.C., P.L., and K.C.H. wrote the paper. All authors reviewed the paper before submission.

## Acknowledgments

The authors thank the Huang and Levin labs for helpful discussions, Paul Lebel for discussions and testing with cell density measurements, and Christine Jacobs-Wagner for generously sharing the µNS strain. Funding was provided by a Stanford Interdisciplinary Graduate Fellowship, an Agilent Graduate Fellowship, and a James McDonnell Postdoctoral Fellowship (to H.S.); a postdoctoral fellowship from the Swiss National Science Foundation under Grant P400PB_180872 (to P.D.O.); a Stanford Graduate Fellowship and National Science Foundation Graduate Research Fellowship (to S.C.); an Arnold O. Beckman Postdoctoral Fellowship (to C.S.W.); the Washington University UStar and SURF programs (to J.K.); NIH R01-GM056836 (to F. C.); NSF CAREER Award MCB-1149328, and the Allen Discovery Center at Stanford on Systems Modeling of Infection (to K.C.H); and NIH R35-GM127331 (to. P.A.L.). K.C.H. is a Chan Zuckerberg Investigator.

## Methods

### Strains and growth media

Cells were initially grown in lysogeny broth (LB) medium or AB+0.2% glucose at 37 °C into log phase. Minimal growth medium made with AB (44) or M9 salts was used for all starvation experiments. Antibiotics were used at the following concentrations: ampicillin, 100 µg/mL; chloramphenicol, 30 µg/mL; rifampicin, 50 µg/mL; cerulenin, 100 µg/mL; A22, 2 µg/mL.

All knockouts were generated in *E. coli* MG1655 cells via transduction from the Keio collection (45).

### Starvation protocol

Cells were first cultured in LB or M9+0.4% glucose and grown to an optical density at 600 nm of 0.1-0.2. Then, 0.8 mL of the culture were transferred to a 1-mL microcentrifuge tube and spun at 8,500*g* for 1 min. The supernatant was immediately removed and 1 mL of AB or M9 medium without carbon was added to the tube to wash the cells. This washing procedure was repeated three times. The medium with the resuspended pellet was placed in a test tube and shaken at 225 rpm at 37 °C for the specified amount of time.

### Single-cell imaging

Agarose pads used for imaging were made by heating a solution containing 1% agarose in the desired growth medium, and then allowing the solution to cool. 1-2 microliters of solution containing starved cells were placed on the pads and allowed to air dry before a coverslip was placed on the agarose pad.

Phase-contrast and fluorescence images were acquired with a Nikon Ti-E inverted microscope (Nikon Instruments) using a 100X (NA 1.40) oil immersion objective and a Neo 5.5 sCMOS camera (Andor Technology). The microscope was outfitted with an active-control environmental chamber for temperature regulation (HaisonTech, Taipei, Taiwan). Images were acquired using µManager v.1.4 (www.micro-manager.org) (46).

*K. pneumoniae* images were acquired on a Nikon Ti-E inverted microscope equipped with a 100X Plan N (NA 1.25) objective (Nikon), SOLA SE Light Engine (Lumencor), heated control chamber (OKO Labs), and ORCA-Flash4.0 sCMOS camera (Hamamatsu Photonics). Fluorescence filter sets were purchased from Chroma Technology Corporation. Nikon Elements software (Nikon Instruments) was used for image acquisition.

### Microfluidics

Overnight cultures were diluted 100-fold into 1 mL of fresh LB and incubated for 2 h with shaking at 37 °C, excepting experiments in minimal medium, for which overnight cultures were diluted into M9+0.4% glucose and incubated for 4 h. Cells were imaged in B04A microfluidic perfusion plates (CellASIC Corp.) and medium was exchanged using the ONIX microfluidic platform (CellASIC Corp.). Plates were loaded with medium pre-warmed to 37 °C. Cells were loaded into the plate, which was incubated at 37 °C, without shaking, for 30 min before imaging.

### Image analysis

The MATLAB (MathWorks) image-processing code *Morphometrics* (47) was used to segment cells and to identify cell outlines from phase-contrast microscopy images. A local coordinate system was generated for each cell outline using a method adapted from *MicrobeTracker* (48). Cell widths were calculated by averaging the distances between contour points perpendicular to the cell midline, excluding contour points within the poles and sites of septation. Cell length was calculated as the length of the midline from pole to pole. See figure legends for the number of cells analyzed (*n*) and error bar definitions.

### Quantitative phase imaging

Cells stained with WGA488 were loaded into the imaging chamber of an ONIX B04A microfluidic chip (CellASIC). Before imaging, Koehler illumination was configured and the peak illumination intensity with 10-ms exposure time was set to the middle of the dynamic range of the Zyla sCMOS 4.2 camera (Andor). µManager v. 1.41 was used to automate acquisition of brightfield *z*-stacks with a step size of 100 nm from ±1 µm around the focal plane (total of 21 imaging planes). Fluorescence images were taken at the focal plane to extract the cell contours. Similar measurements were performed in medium supplemented with 100 mg/mL BSA to calibrate the phase shift equivalent to 100 mg/mL biomass.

For analysis, seven brightfield images at different focal planes (3 slices above and below separated by 200 nm) were used to calculate the phase information using a custom Matlab script implementing a previously published algorithm (17, 18). In brief, this method relates the phase information of the cell to intensity changes along the *z*-direction. Equidistant, out-of-focus images above and below the focal plane are used to estimate intensity changes at various defocus distances. A phase-shift map is reconstructed in a non-linear, iterative fashion to solve the transport-of-intensity equation (17).

A Gaussian peak was fitted to the background of each image and then corrected to be at zero phase shift using an image-wide subtraction of the mean of the peak. Cell contours were segmented using *Morphometrics* v. 1.1 using the fluorescence channel (for shrunken cells, the cell contour includes the region of the enlarged periplasm), and the intensity of pixels inside each cell was extracted to calculate the dry mass density.

### Proteome extraction

Overnight cultures were diluted 1:200 into LB and grown into exponential phase at 37 °C. Before and after 4 h of starvation, ∼10^9^ cells were harvested and washed in cold PBS, then flash-frozen in liquid nitrogen. Two biological replicate samples were collected for each condition. All samples were then thawed and resuspended in 500 µL lysis buffer (6 M urea, 50 mM Tris-base buffer pH 8.1, and 5% sodium dodecyl sulfate) and subjected to 10 min of bead beating. After centrifugation at 10,000*g* for 3 min, the supernatants were reduced with 10 µL of 500 mM dithiothreitol (Millipore Sigma) and alkylated with iodoacetamide (Millipore Sigma). The peptides were washed, digested, and eluted using S-trap tubes (Protifi) following the manufacturer’s protocols, desalted via C18 solid-phase extraction (Sep-Pak Waters), and dried via vacuum centrifugation. Peptide concentration was quantified for normalization using a Nanodrop ND-1000.

### Mass spectrometry proteomic analyses

Peptides were resuspended in 0.2% formic acid to a final concentration of 0.5 µg/µL. Subsequently, 1 µL was loaded onto an in-house laser-pulled 100-µm inner diameter nanospray column packed to ∼22 cm with ReproSil-Pur C18-AQ 3.0 m resin (Dr. Maisch GmbH). Peptides were separated via reverse-phase chromatography on a Dionex Ultimate 3000 HPLC. Buffer A of the mobile phase contained 0.1% formic acid in HPLC-grade water, and buffer B contained 0.1% formic acid in acetonitrile. The HPLC used a two-step linear gradient with 4-25% buffer B for 135 min followed by 25-45% buffer B for 15 min at 0.400 µL/min. Peptides were analyzed on a LTQ Orbitrap Elite mass spectrometer (Thermo Fisher Scientific) in data-dependent mode, with full mass spectrometry scans acquired in the Orbitrap mass analyzer with a resolution of 60,000 and *m/z* range of 340-1,600. The top 20 most-abundant ions with intensity threshold above 500 counts and charge states 2 and above were selected for fragmentation using collision-induced dissociation (CID) with an isolation window of 2 *m*/*z*, normalized collision energy of 35%, activation Q of 0.25, and activation time of 5 ms. The CID fragments were analyzed in the ion trap with rapid scan rate. Dynamic exclusion was enabled with repeat count of 1 and exclusion duration of 20 s. The AGC target was set to 1,000,000 and 50,000 for full FTMS scans and ITMSn scans, respectively. The maximum injection times were set to 250 ms and 100 ms for full FTMS scans and ITMSn scans, respectively.

### Peptide/protein database searching

Mass spectra were searched using Proteome Discoverer 2.2.0.388 using the built-in SEQUEST search algorithm. The Uniprot canonical *E. coli* FASTA database (4350 protein sequences downloaded on 2/9/2020) was used in this search, along with a database containing common preparatory contaminants. The precursor mass range was set to 350-3000 Da, the mass error tolerance was set to 10 ppm, and the fragment mass error tolerance to 0.6 Da. Enzyme specificity was set to trypsin, carbamidomethylation of cysteines (57.021) was set as variable modifications, and oxidation of methionines (+15.995) and acetylation of protein N-terminus (+42.011) were considered as variable modifications. Percolator was used to filter peptides and proteins to a false discovery rate of 1%. Abundance quantification was based on precursor ion peak areas.

### µNS tracking

Analyses were performed as previously described (20). Briefly, cell outlines were extracted using *Morphometrics* (47) from phase contrast images. Local cell coordinates were defined based on the mesh algorithm in *MicrobeTracker* (48), and particle motion was quantified using these local coordinates along the long and short axes of the cell. The fluorescent GFP-µNS particles appeared as diffraction-limited spots, and their positions were determined relative to the cell coordinates by a Gaussian fit in two dimensions.

### Statistical analyses

Two-tailed Student’s *t*-tests were used in Fig. 1E, 2B, and 3B,D,F. Fisher’s exact tests were used in Fig. 1G,H. All *p-*values and sample sizes *n* are included in the corresponding figures and/or figure legends.

## Supplemental Figures

**Figure S1:**
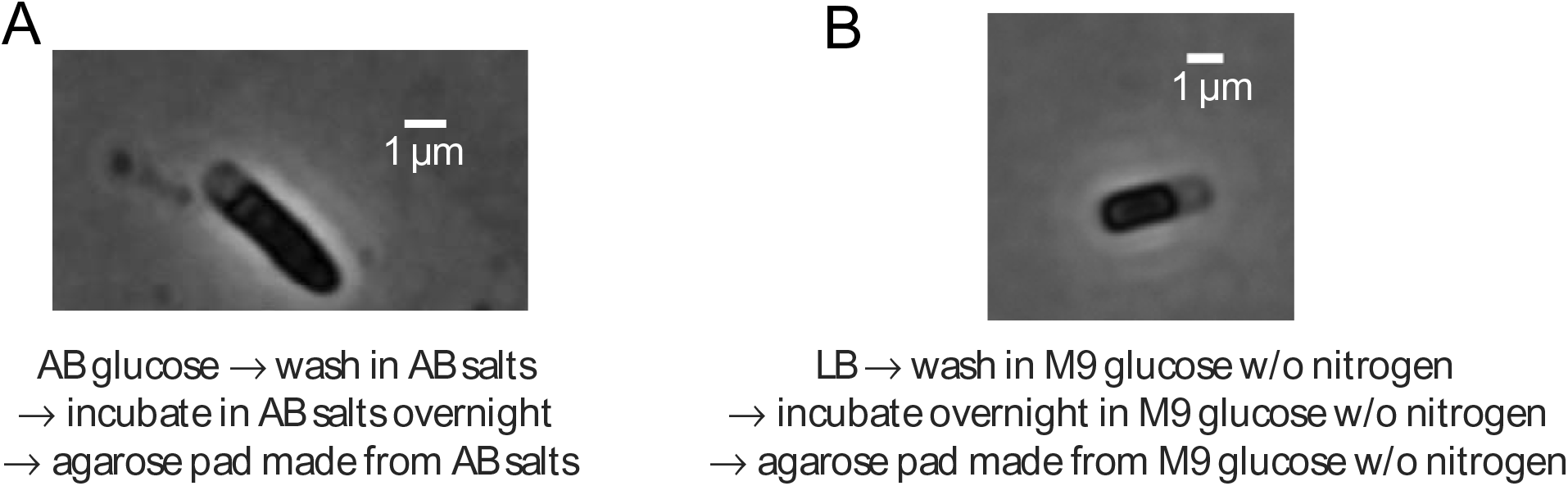
Shrinkage occurs during both carbon and nitrogen starvation. A) Switching from AB glucose to AB salts led to similar shrinkage as switching from LB to M9 salts (Fig. 1B). B) Nitrogen starvation also led to cytoplasmic shrinkage.

**Figure S2:**
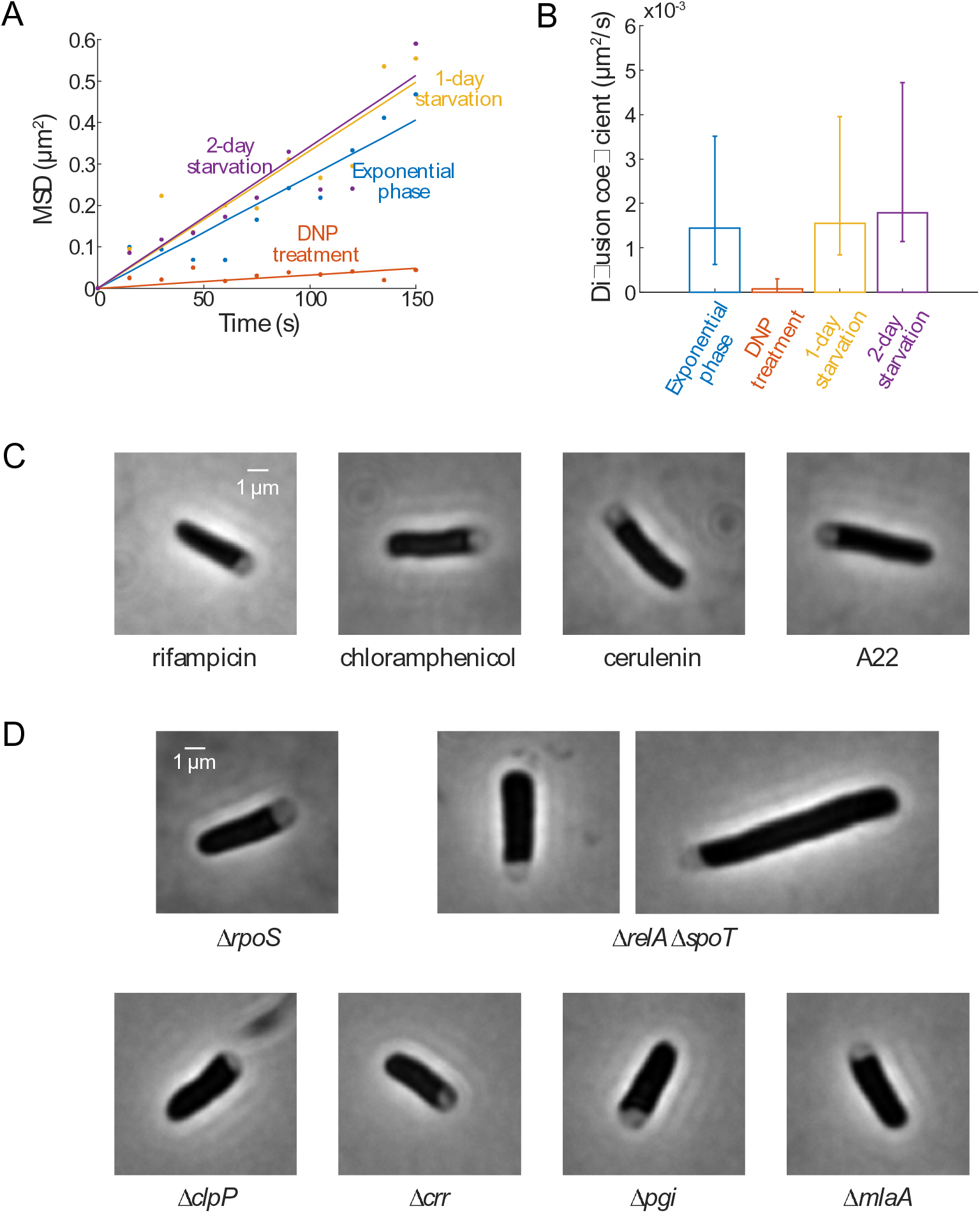
Shrinkage does not affect the diffusion of nanoparticles, and is not disrupted by a wide range of chemical or genetic perturbations. A) Unlike treatment with DNP, carbon starvation did not slow down the diffusion constant D of µNS-GFP particles. Lines are linear fits of the raw data (dots), whose slopes are estimates of 4 times *D. n* > 80 tracks for each condition. B) Carbon starvation did not change the effective diffusion coefficient of µNS-GFP particles. Error bars are 95% confidence intervals. C) Treatment at the time of transfer from LB to M9 salts with antibiotics that disrupt transcription (rifampicin), translation (chloramphenicol), lipid synthesis (cerulenin), or cell-wall synthesis (A22) did not affect shrinkage. D) Shrinkage occurred in mutants that lack stringent-response regulators (Δ*rpoS*, Δ*relA* Δ*spoT*, and Δ*clpP*), key enzymes in glucose processing (Δ*crr* and Δ*pgi*), or components of outer-membrane transport (Δ*mlaA*).

**Figure S3:**
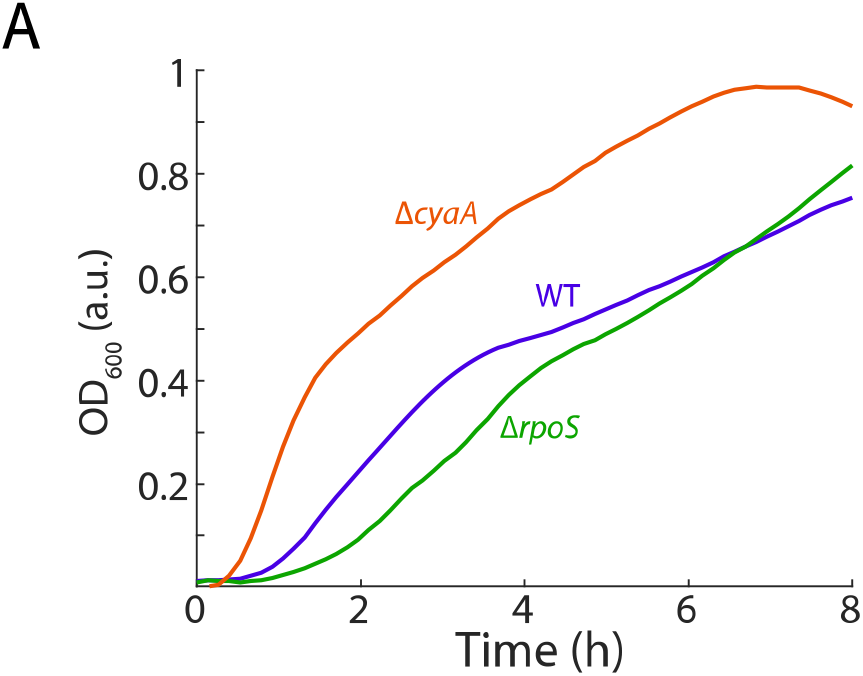
Lag times in deletion mutants. Δ*cyaA* and Δ*rpoS* cells have shorter and longer lag times compared to wild-type cells, which is connected to the different recovery dynamics in Fig. 4B.

**Figure S4:**
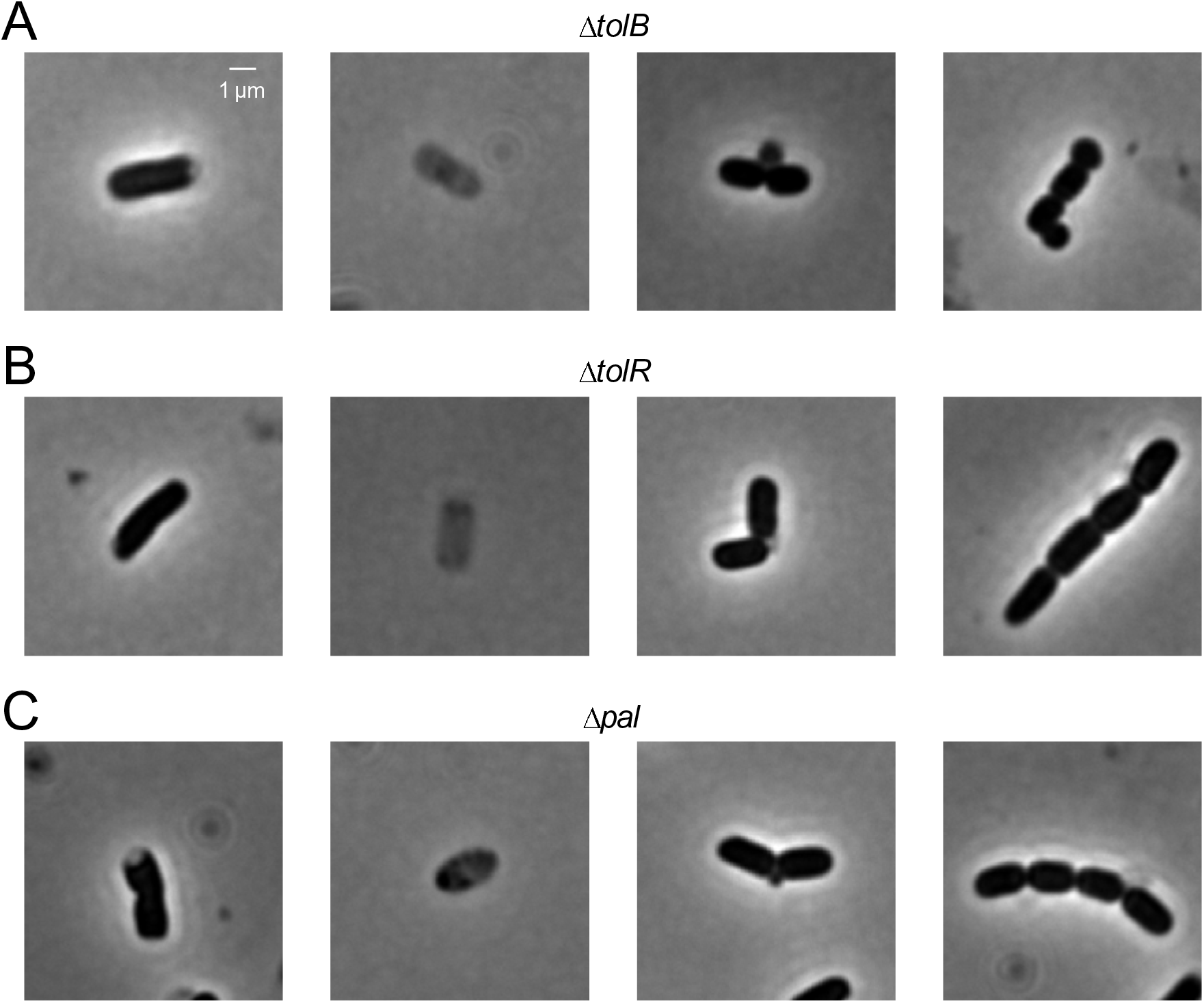
Deletion mutants in the Tol-Pal pathway leads to diverse phenotypes. After 4 h of starvation in M9 salts, Δ*tolB* (A), Δ*tolR* (B), and Δ*pal* (C) cells exhibited diverse phenotypes including shrinkage, lysis, blebbing and chaining, similar as Δ*tolA* (Fig. 6B).

## Supplemental Tables

**Table S1: proteome measurements for log phase and starved cells.**

