## Supplementary material for "Starvation induces shrinkage of the bacterial cytoplasm": Fig. S1

A

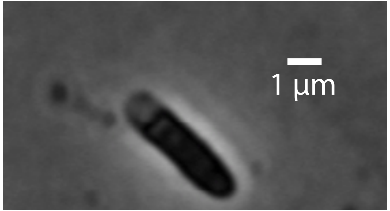


AB glucose → wash in AB salts
→ incubate in AB salts overnight
→ agarose pad made from AB salts

B

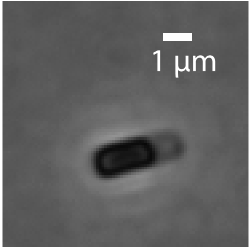


LB → wash in M9 glucose w/o nitrogen
→ incubate overnight in M9 glucose w/o nitrogen
→ agarose pad made from M9 glucose w/o nitrogen
