## Supplementary figures and images for "Starvation induces shrinkage of the bacterial cytoplasm"

### Fig. S2

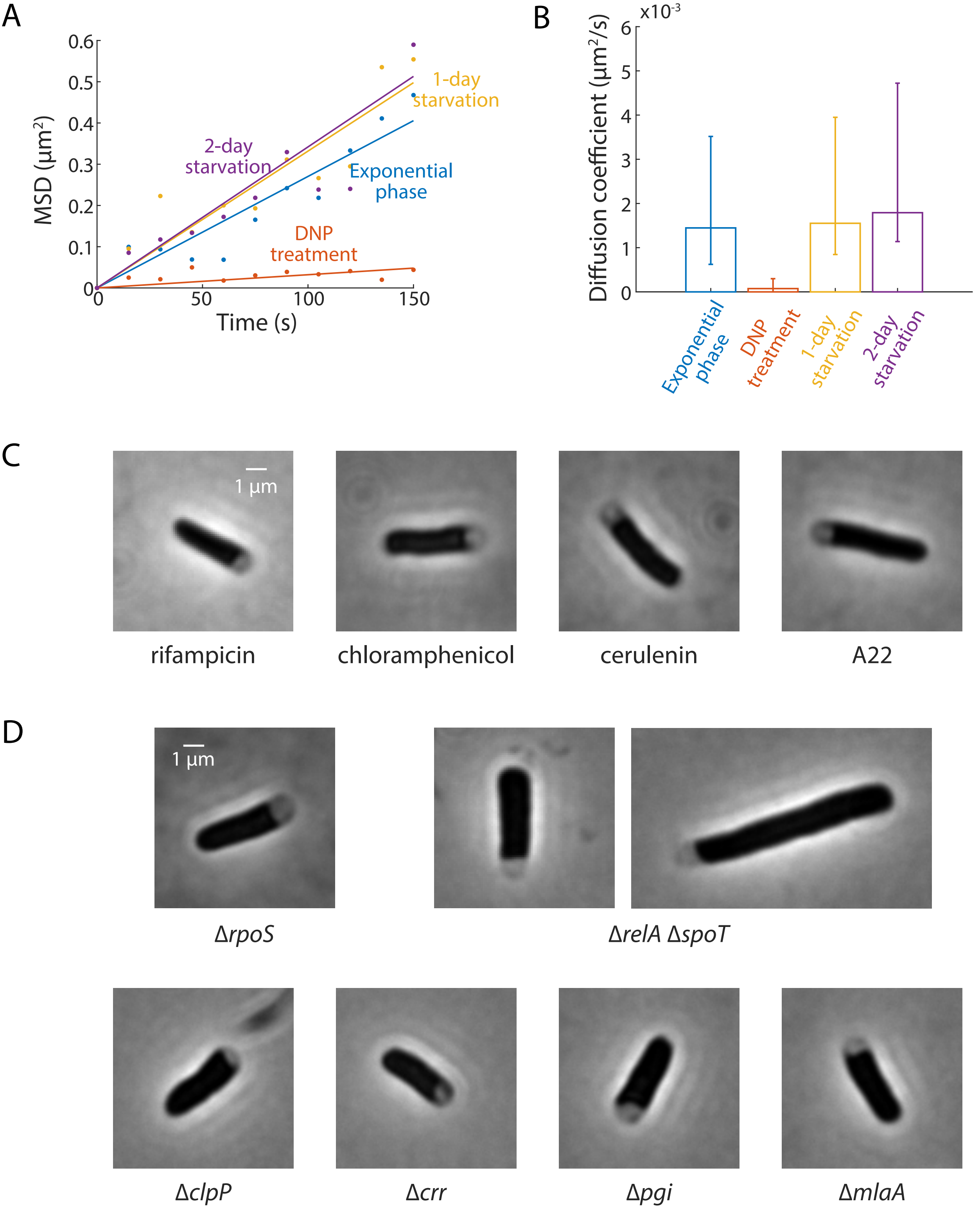

### Fig. S3

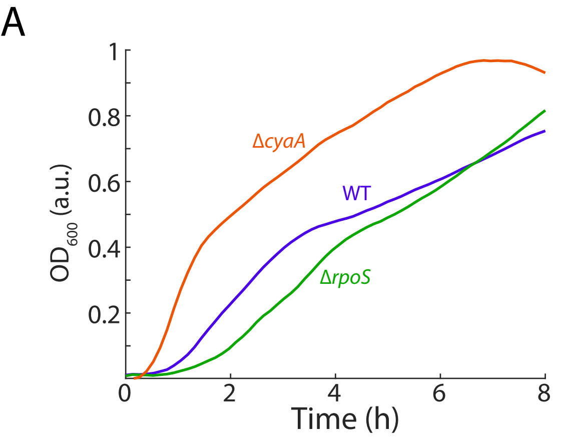

### Fig. S4

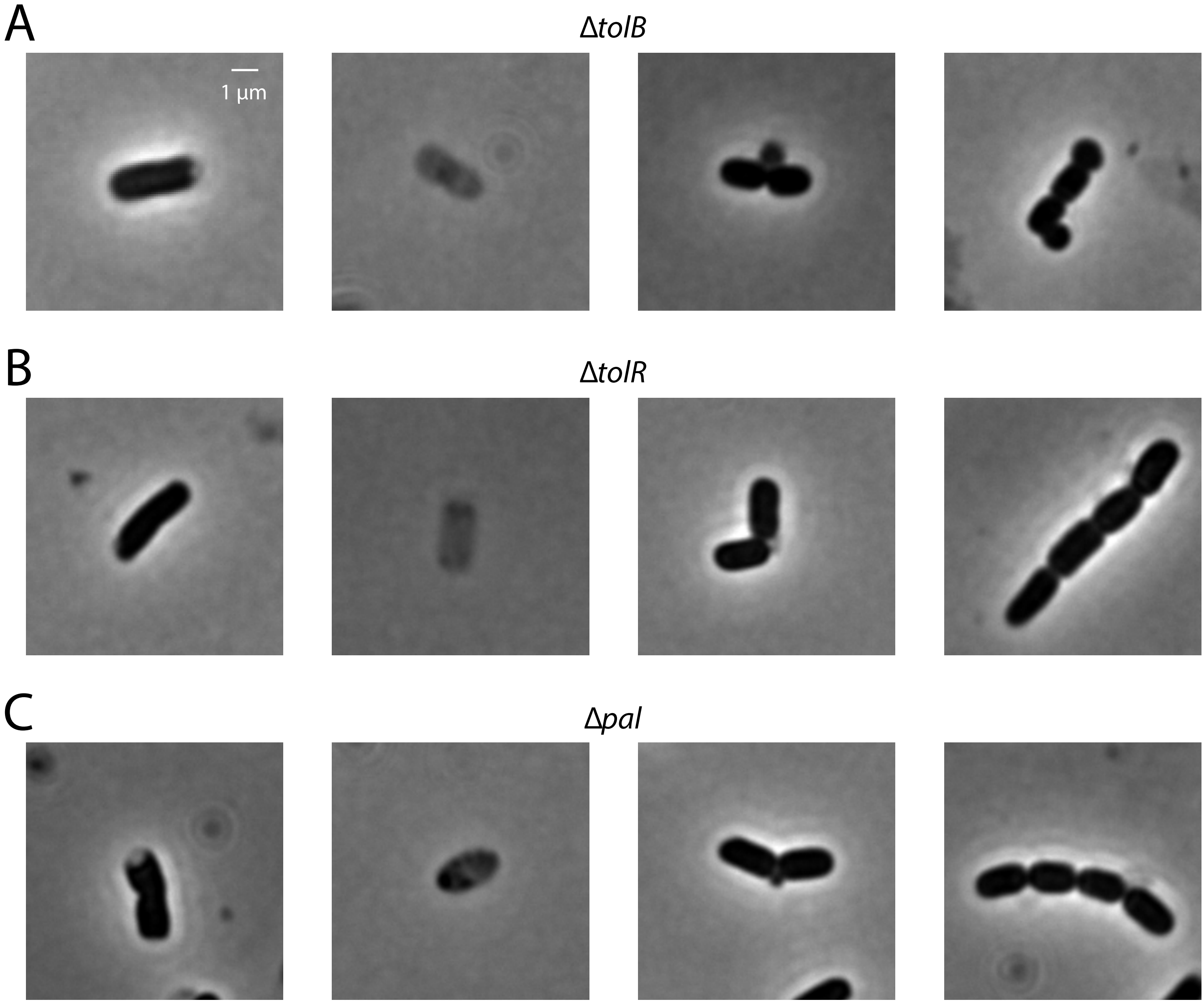
